# Unveiling lipid chemodiversity in root exudates: A comprehensive characterization of the exudate metabo-lipidome in a perennial grass

**DOI:** 10.1101/2024.03.22.586263

**Authors:** Sneha P. Couvillion, Isabella H. Yang, Dylan Hermosillo, Josie Eder, Sheryl Bell, Kirsten S. Hofmockel

**Affiliations:** Earth and Biological Sciences Directorate, Pacific Northwest National Laboratory, Richland, WA, USA

**Keywords:** metabo-lipidomics, root exudates, lipids, mass-spectrometry, tall wheatgrass, metabolite annotation, rhizodeposition, molecular chemodiversity

## Abstract

The rhizosphere, where plant roots meet soil, is a hub of biogeochemical activity with ecosystem impacts on carbon stocks. Root derived carbon has been found to contribute more to soil carbon stocks than aboveground litter. Nonetheless, the molecular chemodiversity of root exudates remains poorly understood due to limited characterization and annotation. Here our goal was to discover the molecular chemodiversity of metabolites and lipids in root exudates to advance our understanding of plant root inputs belowground. We worked with mature, field-grown tall wheatgrass (*Thinopyrum ponticum*) and optimized exudate collection protocols to enable the capture of non-polar lipids in addition to polar and semi-polar metabolites. Rates of carbon input via hydrophobic exudates were approximately double that of aqueous exudates and carbon/nitrogen ratios were markedly higher in hydrophobic compared to aqueous exudates, emphasizing the importance of lipids, due to their high carbon content. To maximize molecular coverage of exudate chemodiversity, we used liquid chromatography coupled tandem mass-spectrometry for paired untargeted metabolomics and lipidomics or ‘metabo-lipidomics’. We substantially increased the characterization of exudate chemodiversity by employing both tandem mass spectral library searching and deep learning-based chemical class assignment. Notably, in this unprecedented characterization of intact lipids in root exudates, we discovered a diverse variety of lipids, including substantial levels of triacylglycerols (∼19 μg/g fresh root per min), fatty acyls, sphingolipids, sterol lipids, and glycerophospholipids. Comparison of the root exudate and tissue lipidomes revealed minimum glycerophospholipids in exudates, suggesting the exudate protocol did not extract lipids from root cell membranes.

## Introduction

### Root Exudates: Major Carbon Inputs and Regulators of Soil Organic Matter (SOM) Dynamics

Current estimates of direct plant contributions to SOM pools such as mineral associated organic matter (MAOM) are likely undervalued given the reliance on aboveground plant tissue proxies for determining plant versus microbial origin (Whalen *et al*., 2022). This is in part due to the challenging nature of quantifying rates, chemical composition and ecological significance of root derived rhizodeposits including exudates (Phillips *et al*., 2008). The roles of small molecule metabolites, especially lipids, in shaping interkingdom interactions and microbial processes that control cycling and long-term storage of carbon are poorly understood. Metabolites and lipids play diverse roles in regulatory, metabolic, and signaling networks among microbes (Whiteley *et al*., 2017, Khalid & Keller, 2021) and in microbe-plant interactions (Chagas *et al*., 2018) and are sensitive indicators of organismal phenotypes and community metaphenomes (Jansson & Hofmockel, 2018). However, challenges in identifying key molecules and interpreting their functions are critical frontiers in soil ecosystem research (Brown *et al*., 2024).

Plants carry out the necessary first step of siphoning atmospheric carbon into soil through photosynthesis. It is estimated that 5-40% of photosynthetically fixed carbon enters soil through root exudation (Simon & Haichar, 2019, BADRI & VIVANCO, 2009). Exudates regulate SOM dynamics by contributing to SOM stabilization and turnover. Small molecules in root exudates, such as lipids, function as both signaling compounds and microbial nutrients that shape the assembly and activity of microbial communities (Huang *et al*., 2019, Zhalnina *et al*., 2018, Mark *et al*., 2005, Hugoni *et al*., 2018, Sasse *et al*., 2018). They influence the colonization, growth, and activity of key microbial members, including mycorrhizal fungi (Tian *et al*., 2021, Elias & Safir, 1987), impact soil physical structure (Naveed *et al*., 2017), induce soil respiration (de Vries *et al*., 2019), and can cause priming, decomposition, and mineralization of SOM. These small molecules may regulate microbial metabolism, undergo assimilation into microbial biomass, transformation by extracellular enzymes, or association with soil minerals, ultimately integrating into SOM. Research on priming (Fontaine *et al*., 2007, Chen *et al*., 2019) and factors controlling the formation of stable SOM (Liang *et al*., 2023, Kallenbach *et al*., 2016, Rempfert *et al*., 2024) have predominantly used single or simplistic plant carbon input surrogates such as glucose, cellulose and cellobiose. However, this approach overlooks the realistic chemodiversity present in exudates. By relying on these surrogates, the studies tend to simplify plant inputs to carbon sources alone, neglecting the diverse metabolic, regulatory, and signaling roles played by the small molecules present in exudates. Different chemical classes of root exudate surrogates induce distinctly different magnitudes of priming and these effects are likely non-additive in complex mixtures, emphasizing the complexity of their effects on SOM stocks (Yan *et al*., 2023) and the need to consider root exudate molecular complexity.

### Capturing Exudate Chemodiversity using Metabo-Lipidomics: Navigating Beyond just Sugars and Organic Acids into the Uncharted Territory of Lipids

Recent studies using liquid chromatography coupled with mass spectrometry (LC-MS/MS) for untargeted metabolomics have revealed a broad spectrum of polar and semi-polar small molecules in exudates (McLaughlin *et al*., 2023, Dietz *et al*., 2019). While progress has been achieved in characterizing these metabolites, the comprehensive identification of the numerous features detected by mass spectrometry remains a significant bottleneck in metabolomics, with only a small fraction of features being currently identified (da Silva *et al*., 2015, Giera *et al*., 2022, Shen *et al*., 2023). Importantly, the exploration of lipids, a crucial group of non-polar metabolites, is notably absent in root exudate studies. Their distinctive hydrophobic properties present challenges in detection and analysis using conventional methods for exudate collection and analysis. Our recent work demonstrates the value of intact lipids in soil ecosystems as sensitive indicators of environmental stress response and substrate availability and highlights their great potential for interrogating interkingdom interactions and resource sharing (Couvillion *et al*., 2023, Naasko *et al*., 2023).

In this study, we hypothesized that plant roots exude a diverse array of intact lipids, potentially serving as microbial energy sources, chemotactic signals, antimicrobial compounds, and metabolic regulators in the rhizosphere. Accurately assessing plant exudate profiles, which are significantly influenced by growth stage and growth conditions, requires collecting root exudates from mature plants in a way that realistically recapitulates natural field conditions (Dietz et al., 2019, Williams *et al*., 2021). This was highlighted by a recent study which found higher total carbon in hydroponic exudates, whereas field-grown plant exudates contained more secondary metabolites-suggesting hydroponic root exudates may not fully capture the exudate composition and chemodiversity of field environments (Heuermann *et al*., 2023). We chose to work with mature, field grown tall wheatgrass plants (*Thinopyrum ponticum*, cultivar: Alkar) to maximize translation to perennial plants in the field. To collect the full diversity of small molecules, we optimized exudate collection protocols to enable the capture of non-polar (hydrophobic) lipids in addition to polar and semi-polar (hydrophilic) metabolites. We found that C and N input rates in hydrophilic and hydrophobic exudates were 38.41 ± 5.93 and 81.52 ± 13.81 μg C g^-1^ fresh root mass min^-1^, respectively, and 2.68± 0.42 and 0.20 ± .04 μg N g^-1^ fresh root mass min^-1^, respectively (**Fig. 1**). The C/N of hydrophilic exudates was far lower than that of hydrophobic exudates (14.40 ± 0.58 vs 459 ± 90), which is consistent with high C content in lipids, and points to potential differences in metabolite and lipid contributions to SOM dynamics. To explore the molecular diversity of root exudates, We used liquid chromatography coupled tandem mass-spectrometry (LC-MS/MS) for paired untargeted metabolomics and lipidomics or ‘metabo-lipidomics’ (**Fig. 2A**); we employed three liquid chromatography column chemistries and analyzed samples in positive and negative ionization modes to maximize coverage of the metabo-lipidome. The results from the metabolomics and lipidomics analyses will be discussed separately.

**Figure 1:**
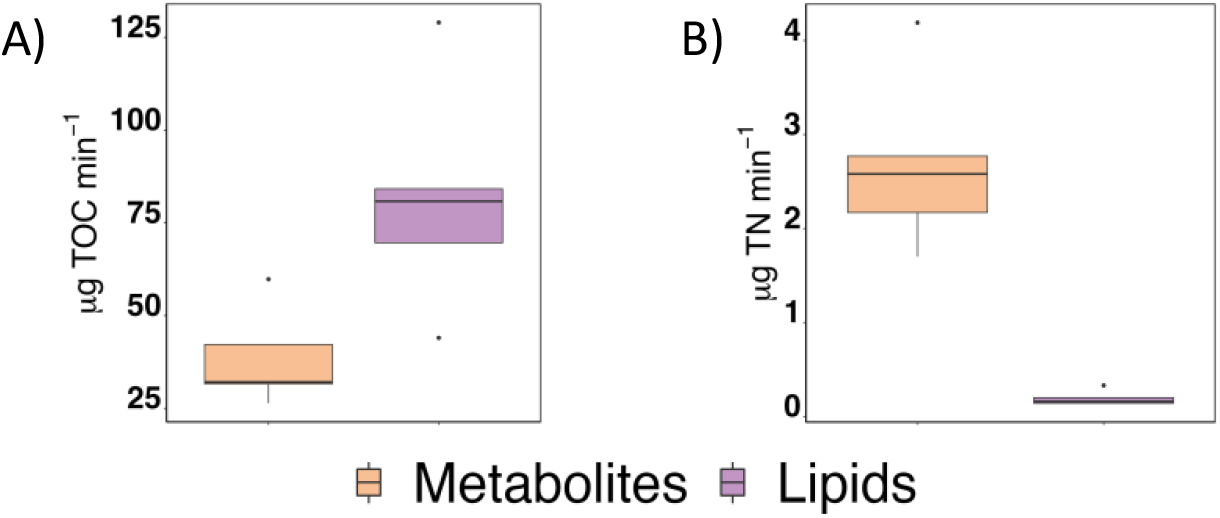
A) Total organic carbon (TOC) and B) total nitrogen (TN) input rates in aqueous metabolite (orange) and hydrophobic lipid (lilac) root exudate fractions, normalized to root mass

**Figure 2:**
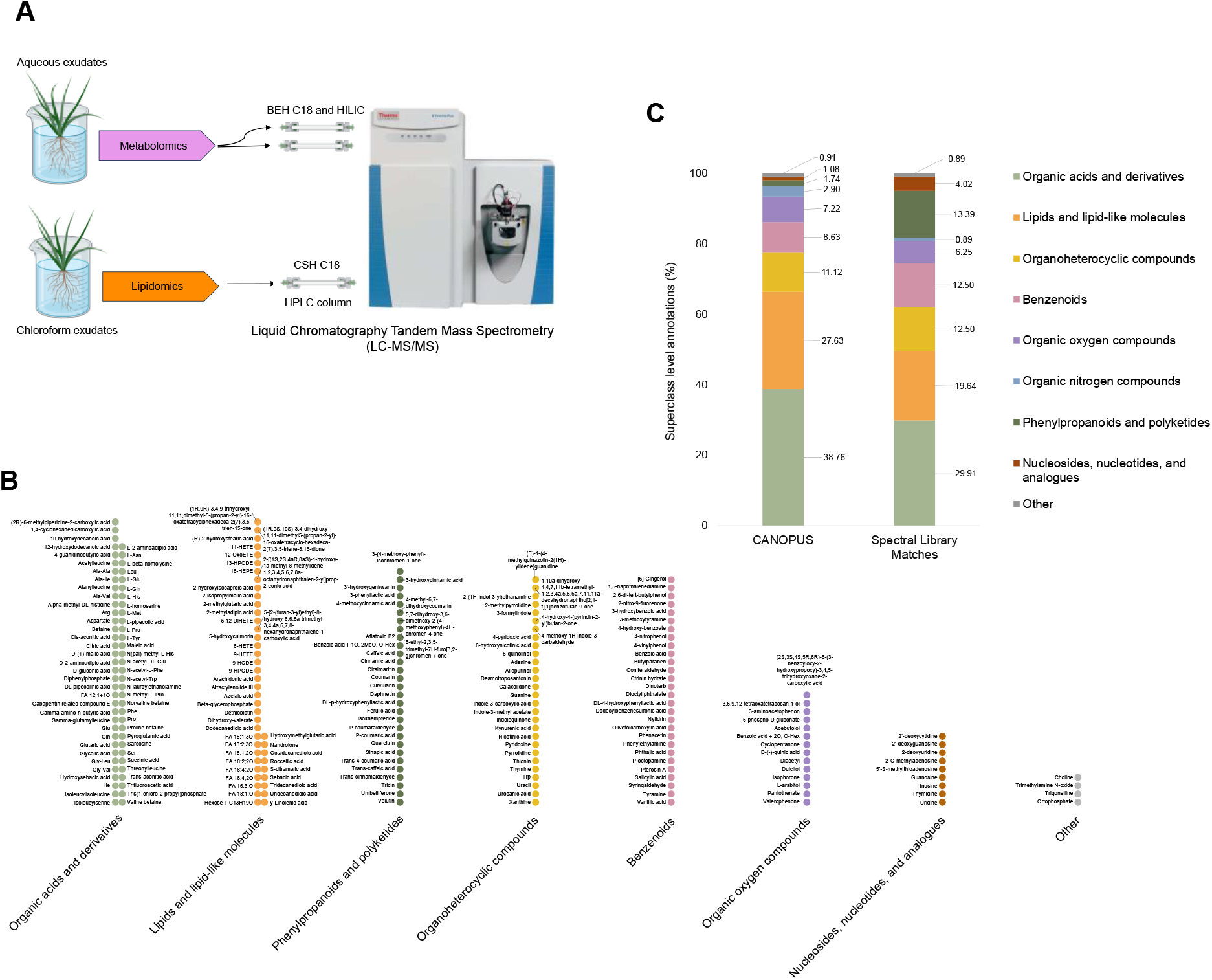
A) Untargeted LC-MS/MS based metabo-lipidomics workflow; B) Metabolites annotated using spectral library searching, colored and listed by superclass C) Percent distribution of superclass level classification of metabolomics features annotated using CANOPUS and spectral library searching (SLS)

### Tackling the Unknowns: Enhancing Metabolite Annotation and Classification by Combining Spectral Library Searching with CANOPUS

Metabolite identification in mass-spectrometry analyses poses a major challenge in metabolomics research (Monge *et al*., 2019). To address this, our study combines two strategies for improved feature annotation. We employed spectral library searching (SLS) to assign annotations to 224 unique metabolites (Schymanski confidence level 2A (Schymanski *et al*., 2014), **Fig 2B**), using in-house and publicly available MS/MS databases. These annotations were established through accurate mass and tandem mass spectra matching, with additional consideration of retention time when there was a match to the in-house library. Using the SLS approach we identified 67 metabolites belonging to the ‘Organic acids and derivatives’ superclass, primarily comprising amino acids and analogues, and dipeptides. The ‘Lipids and lipid-like molecules’ superclass ranked second, encompassing 44 metabolites, predominantly composed of fatty acids, derivatives, and prenol lipids. The ‘Phenylpropanoids and polyketides’ superclass contained 30 metabolites, including cinnamic acids and derivatives, flavonoids, coumarins, phenylpropanoic acids, and cinnamaldehydes. Similarly, the superclasses ‘Organoheterocyclic compounds’ (indoles, pyridines, quinolines, and derivatives) and ‘Benzenoids’ (benzene derivatives, phenols) each comprised 28 metabolites. Finally, the superclasses ‘Organic Oxygen Compounds’ and ‘Nucleosides, nucleotides, and analogues’ contained 14 and 9 metabolites, respectively.

These finding show that field grown tall wheatgrass root exudates contain a rich diversity of metabolites spanning various superclasses, some of which are recognized for their roles in mediating stress responses, plant-microbe interactions, nutrient cycling, and defense mechanisms. In addition to primary metabolites, we identified a diverse array of secondary metabolites including benzoic acid derivatives, prenol lipids, indoles and derivatives, cinnamic acids, coumarins and flavonoids. Metabolites belonging to these subclasses have been shown to have a role in plant defense, stress response, microbial recruitment, and signaling pathways (Schenkel *et al*., 2015, Van Poecke *et al*., 2001, Massalha *et al*., 2017, Tian et al., 2021, Zhang *et al*., 2009, Hassan & Mathesius, 2012). Among the 224 metabolites that we identified through spectral library matching, we found that 43 have been previously reported (McLaughlin et al., 2023, Zhalnina et al., 2018, Strehmel *et al*., 2014) in root exudate characterizations of other plants. Notably, 30 of these previously reported metabolites were recently characterized as part of the core exudate metabolome (McLaughlin et al., 2023) in three model plants—*Arabidopsis thaliana, Brachypodium distachyon*, and *Medicago truncatula*—constituting 40% of the 75-molecule core exudate metabolome. The core metabolites prominently featured nucleobases (adenine, guanine, thymine, uracil) and organic acids, particularly amino acids (asparagine, glutamic acid, glutamine, histidine, leucine, proline, tyrosine), indicating a commonality of these small molecules in exudates across diverse plant species.

While database-dependent methods such as SLS, offer valuable insights into chemical structures, their identification rate is constrained by the library’s scope, leaving a substantial majority-often ∼90%-of features unannotated (da Silva et al., 2015, Shen et al., 2023). This is a central challenge in the field of metabolomics that needs to be addressed in order to obtain ecological insights from molecular level data (Peters *et al*., 2018). To maximize the biological information obtained from the unannotated features which are typically discarded from downstream analyses, we complemented SLS with CANOPUS (Dührkop *et al*., 2021), a deep learning-based class assignment and ontology prediction tool. This enabled the prediction of chemical classifications for 1559 detected, high quality features. It’s important to note that due to the higher masses of many lipids, CANOPUS analysis was exclusively applied to the metabolomics data. CANOPUS predicted classifications for 80% of these features spanning 12 superclasses (1205 features classified), 56 classes (1084 features), 85 subclasses (822 features), and 60 level 5 classifications (588 features). Approximately 1250 features received at least one level of classification, substantially increasing our characterization of the chemodiversity of plant exudates. The SLS and CANOPUS analyses uses two separate software (MS-DIAL and MZmine) for feature detection and alignment; it wasn’t possible to match features across the analyses. Nonetheless, the combined approach resulted in expanding current knowledge of root exudate chemodiversity.

The superclass-level distribution of annotations from spectral library search (SLS; 224 features) and CANOPUS (1205 features) exhibited similar trends (**Fig 2C**). ‘Organic acids and derivatives’ predominated in both approaches (SLS 30%, CANOPUS 39%), comprising mainly amino acids, derivatives, and peptides. Following closely was the ‘Lipid and lipid-like molecules’ superclass (SLS 20%, CANOPUS 28%), where fatty acids and conjugates were prominent. ‘Organoheterocyclic compounds’ (SLS 12.5%, CANOPUS 11%), ‘Benzenoids’ (SLS 12.5%, CANOPUS 9%), ‘Organic oxygen compounds’ (SLS 6%, CANOPUS 7%), including carbohydrates and carbonyl compounds, ‘Organic nitrogen compounds’ (SLS 1%, CANOPUS 3%), ‘Phenylpropanoids and polyketides’ (SLS 13%, CANOPUS 2%), and ‘Nucleosides, nucleotides, and analogues’ (SLS 4%, CANOPUS 1%) followed.

### Discovering the Diversity of Lipids in Root Exudates

In the untargeted lipidomics data, 7864 features with MS/MS spectra were detected in positive and negative ionization modes combined. Of these, a mere 289 (∼4%) lipid features (including isomers) comprising 267 unique InChIKeys(Heller *et al*., 2015) were annotated using MS/MS database matching (Schymanski identification confidence level 2A (Schymanski et al., 2014)) with MS-DIAL. Lipids from the glycerolipid and fatty acyl categories were particularly prevalent, followed by sphingolipids, sterol lipids, and glycerophospholipids [**Fig. 3A**]. Known functions of these lipids include diverse roles in membrane structure and stability, energy storage, cell signaling, and responses to stress.

**Figure 3:**
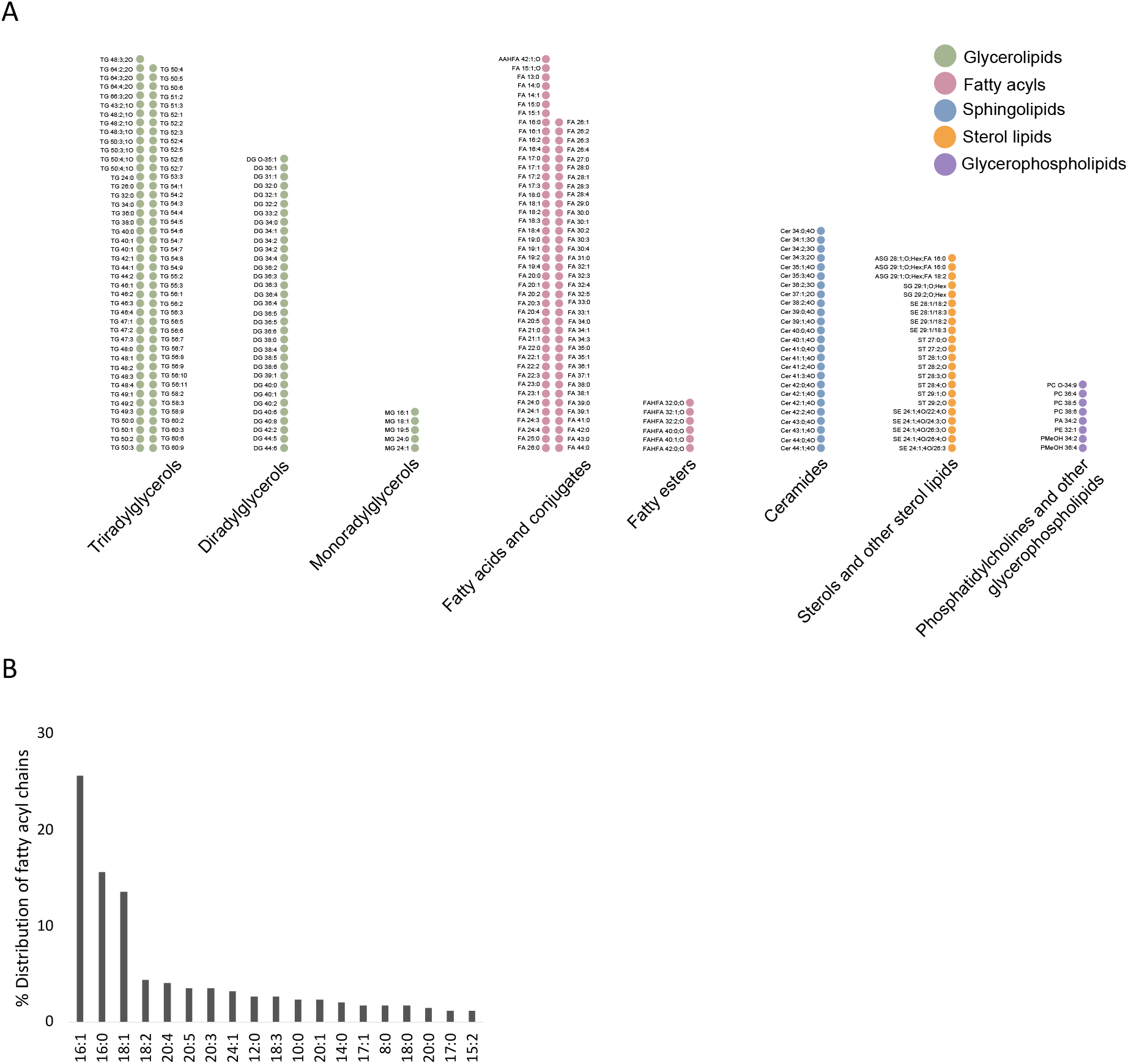
A) Level 2A feature annotations in untargeted lipidomics data; grouped and colored by lipid main class B) Distribution of fatty acid residues in glycerolipids

In hydrophobic root exudates of mature tall wheatgrass, we found that glycerolipids, particularly triacylglycerols (TG), represent a dominant fraction, comprising 47% of the lipidome. Subclass analysis reveals the prevalence of triradylglycerols (TG), diradylglycerols (DG), and monoradylglycerols (MG) at 33%, 12%, and 2% of annotated lipids, respectively. We examined the distribution of fatty acids within the glycerolipids to identify the top three predominant fatty acids—16:1, 16:0, and 18:1—followed by 18:2, 20:4, 20:5, 20:3, and 24:1, shaping the composition of TG, DG, and MG in root exudates (**Fig 3B**). To approximately quantify glycerolipids in exudates, we spiked a deuterated standard for each subclass. During a 5-minute hydrophobic exudate collection, roots exuded 2 μg of MG, 17.2 μg of DG, and 95.1 μg of TG per gram of fresh root mass. Extrapolating this rate suggests a noteworthy daily exudation of approximately 806 mg of TGs per plant. This calculation represents one of the first approximations of compound-specific exudation rates for mature perennial plants-a central, but missing value that is necessary for quantifying exudate effects on SOM dynamics (citation).

Few studies have explored TG lipid secretion in higher plants. A recent study (Tatsumi *et al*., 2023) revealed significant TG secretion from both cultured cells and hairy roots of *Lithospermum erythrorhizon*, with cultured cells secreting about 30% of total TG produced. The authors also observed TG secretion in *N. tabacum* cultured cells and intact plant roots, indicating its occurrence across plant species, consistent with our tall wheatgrass findings. Interestingly, TGs were found to act as carrier lipids, forming lipid droplets that transport other hydrophobic metabolites extracellularly. This confirms that the chemodiversity of root exudate lipids found in this study is not merely an experimental artifact. While the molecular mechanisms behind TG secretion remain unknown, bayberry (*Myrica pensylvanica*) fruits accumulate TG-rich surface wax through repurposed cutin synthesis genes (Simpson *et al*., 2016). Diurnal variations in exudation rates (McLaughlin et al., 2023) and the stress induced by the collection protocol may impact TG metabolism and exudation rates, given TGs’ role in oxidative stress protection (Fan *et al*., 2017).

Further investigation into these influences is warranted. In addition to acting as carriers lipids (Tatsumi et al., 2023) we propose that these lipids may influence microbial dynamics, serving as a nutrient source, fostering symbiotic relationships, and impacting the formation and stabilization of water-stable aggregates. Microbial breakdown of lipids such as triacylglycerols in soil would require extracellular enzymes such as esterases and lipases and studies suggest that these lipids may persist longer than simple sugars in soil (Soliman & Radwan, 1981, Hita *et al*., 1996).

Amongst the fatty acyl category, 30% of the identified lipids belonged to the subclass of free fatty acids (FA). Fatty acids and derivatives are known to carry out important functions such as stimulating P-solubilizing activity (Pantigoso *et al*., 2023) and denitrification activity (Lu *et al*., 2014) in rhizosphere bacteria as well as sustaining AMF germination and sporulation (Kameoka & Gutjahr, 2022). However, given the sparse tandem mass spectral fragmentation of FAs, future work needs to confirm the identity of FAs using standards for retention time confirmation and/or gas chromatography coupled with mass spectrometry. Aside from FAs, the remaining fatty acyls consisted of six fatty acyl esters of hydroxy fatty acids (FAHFAs) and one acyl alpha-hydroxy fatty acid (AAHFA). FAHFAs are a recently discovered class of bioactive lipids found in plants, yeast, and mammals, some of which exhibit anti-inflammatory and anti-diabetic activity (Kolar *et al*., 2019, Yore *et al*., 2014). The role of these lipids in root exudates is still unknown.

Sphingolipids—specifically ceramides—made up 9% of the annotated lipids [**Fig. 2A**]. Ceramides serve many functions in plants, including cell signaling, stress response, and membrane integrity. The biophysical properties and functions of ceramides can vary depending on the constituent sphingoid base and fatty acid. The ceramides we detected were largely composed of trihydroxy long chain bases (LCB) 18:0 and 18:1 (48% and 16% of the ceramides respectively) and α-hydroxy fatty acids varying in length from 16 to 26 carbons which are typical in plants (Lynch & Dunn, 2004). Isolated LCBs have been shown to inhibit the growth of plant-associated fungi and bacteria *(Glenz et al*., *2022)*. Ceramides in other host-microbe ecosystems such as human skin are known to inhibit pathogenic biofilm formation (Cleary *et al*., 2018). The functions of ceramides in root exudates are yet to be elucidated, but their potential roles in regulating the growth of plant-associated microbes and biofilms warrant further investigation given their abundance in root exudates.

Phytosterols, accounting for 8% of identified lipids, play key roles in plant-microbe interactions and plant health. We identified stigmasterol, a plant sterol with demonstrated potential for improving nitrogen use efficiency in the rhizosphere. A recent study revealed that a denitrifying bacterial inoculant strain could influence stigmasterol concentrations in duckweed root exudates leading to increased nitrite reductase activity, biofilm formation, and promotion of plant growth (Lu *et al*., 2021). Only eight glycerophospholipids (3%) were identified and these included phosphatidylcholines and phosphatidylethanolamine which are abundant phospholipids in plant cell membranes. The low number of phospholipids detected suggests that the hydrophobic exudate collection protocol used in this study effectively captured lipids arising from root exudates rather than from root cell membranes or surface attached microbial cells.

### Comparison of the Root Exudate Lipidome with the Root Tissue Lipidome

We used untargeted LC-MS/MS lipidomics to characterize lipids in root tissue and compared root tissue and exudate lipid composition. In root tissue, 539 lipid features (including isomers) comprising 503 unique InChIKeys were annotated using MS/MS database matching (Schymanski identification confidence level 2A) with MS-DIAL. Lipids from the glycerolipid, glycerophospholipid, fatty acyl, sphingolipid, and sterol lipid categories were detected [**Fig. 4A**]. Upon comparing lipids detected in root exudates and root tissue, we found that amongst a total of 594 lipids, 327 were only detected in roots, 91 were only detected in root exudates and 176 were detected in both roots and root exudates [**Fig. 4B**]. Although it is unclear why certain molecules were detected in exudates but not in root tissue, this has been observed previously by others (McLaughlin et al., 2023). A breakdown of lipids by category, unique to roots or exudates or those detected in both is shown in **Fig. 4C**. While only 8 glycerophospholipids were detected in exudates, 102 glycerophospholipids were detected in root tissue. We examined the distribution of fatty acyl chains in the triacylglycerols and found that in TGs detected only in the roots, 18:2 was predominant (51% of all fatty acyl chains) followed by 18:1 (22%) and 18:3 (22%). Interestingly, in TGs detected only in the exudates, 20:3 constituted 9% of the fatty acyl chains,

**Figure 4:**
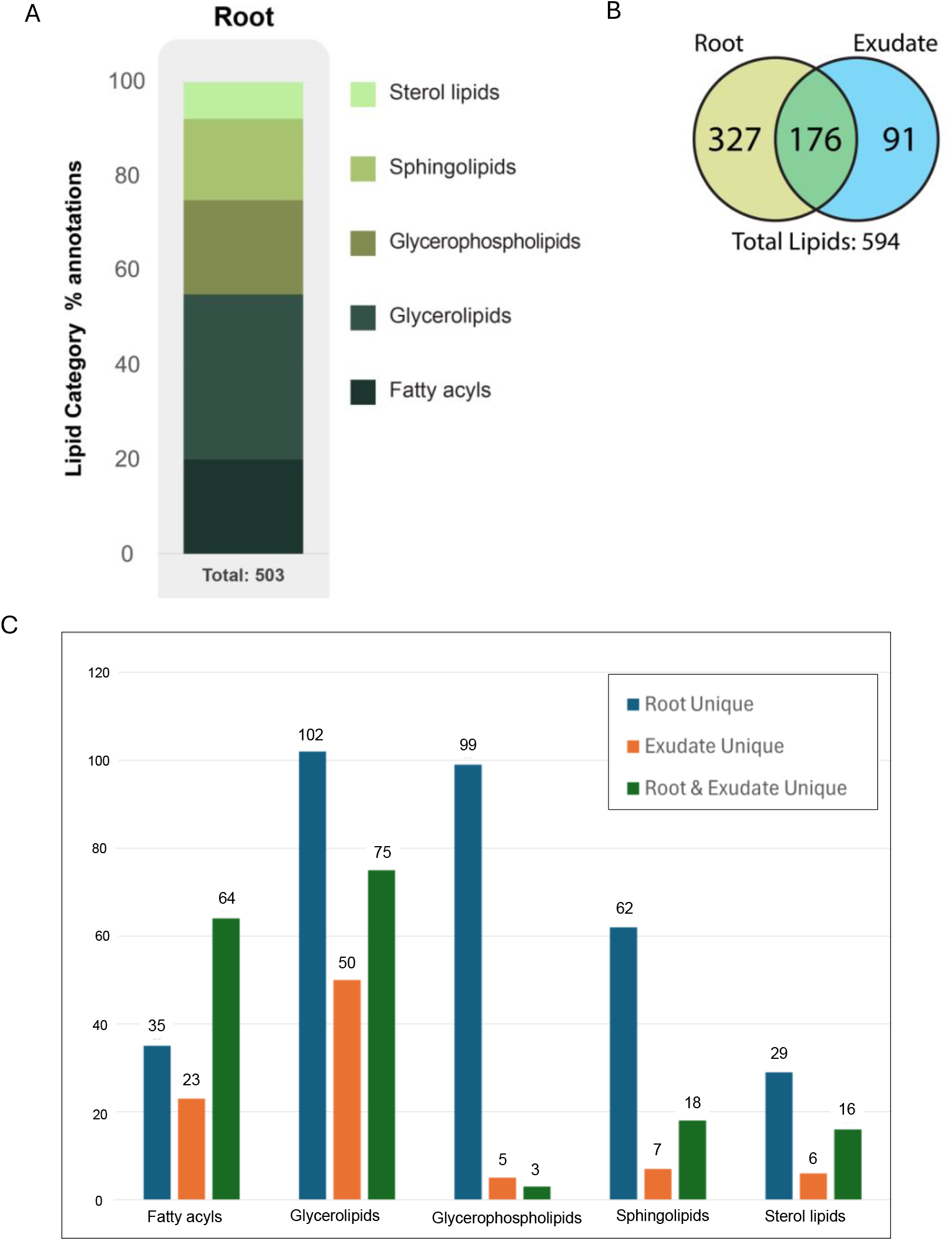
A) Percent distribution of lipid categories of lipids identified in roots B) Venn diagram of unique and overlapping lipids identified in root tissue and exudates C) Breakdown of number of unique and overlapping lipids identified in roots and exudates by lipid category while it was not present in any of the TGs detected in only the roots. Detailed lists of all identified lipids and metabolites in this study can be found in the **Supplementary Files**.

### Conclusions and Future Directions

A comprehensive characterization of small molecules-including lipids-in root exudates is a critical first step for advancing our understanding of the functions and fate of rhizodeposits in terrestrial ecosystems. Using mature, field-grown tall wheatgrass plants as a case study, we demonstrate the first comprehensive characterization of the root exudate metabo-lipidome. We found that C input rates in hydrophobic exudates were around double that of aqueous exudates and C/N ratios were markedly higher in hydrophobic exudates when compared to aqueous (459 ± 90 vs 14.40 ± 0.58). Despite the caveat that there may be variations in exudation in undisturbed soil environments and our collection methods that use water and chloroform, these findings emphasize the need to pay closer attention to the lipid fraction of root exudates and highlight the need to investigate the impact of exudate lipids on microbial physiology, carbon use efficiency, and SOM formation and turnover. In an unprecedented characterization of intact lipids in root exudates, our results reveal the presence of a variety of lipids with diverse functions, with substantial levels of triacylglycerols which previously were not known to be exuded by roots. The combined use of SLS and CANOPUS strategies for metabolite identification, not only enabled high confidence annotations via spectral library searches but also facilitated a broader assignment of chemical class information across detected features. This dual strategy, leveraging the strengths of both methods, enhanced the depth and coverage of metabolite identification, contributing to a more thorough understanding of the exudate metabolome. Overall, with 513 metabolite and lipid identifications and chemical classification of 1250 features (∼80% of all high-quality features detected), this work represents the most comprehensive molecular characterization of root exudates so far.

Achieving sustainable and climate-resilient ecosystems requires an understanding of the impact of root exudates on microbial community functions and soil carbon dynamics. The findings in this work will be critical in resolving current uncertainties in quantifying plant contributions to SOM formation and stabilization. Future studies need to quantify the impacts of environmental factors (climate, soil properties) on the exudate metabo-lipidome, and evaluate the functional consequences of various small molecules on microbial metabolism and activity. Leveraging the findings in this work, targeted incubations with exudate metabolites and lipids can be designed to understand the effect of these molecules on microbial metabolism and their interactions with minerals. The impact and contributions of root endophytes and symbionts such as arbuscular mycorrhizal fungi on exudate chemistry is not known and warrants interrogation. Investigations on priming can incorporate increasingly complex mixtures or natural exudates to understand microbial community responses in realistic conditions. Comprehensive molecular mapping of the soil metabo-lipidome, by systematic characterization of small molecule metabolites and lipids from plant and microbial sources, and their cross-feeding between organisms, is essential for understanding interactions and metabolic pathways that drive ecological functions such as SOM formation and nutrient cycling.

As we navigate the open frontier of exploring the functions of lipids in root exudates and soil ecosystems, particularly their potential to enhance plant health, long-term storage of soil carbon and ecosystem resilience, the identification of key bioactive exudate metabolites and lipids becomes paramount. This knowledge can inform breeding of desired plant traits to facilitate plant-microbe partnerships that can enhance plant productivity and stress tolerance, all while promoting soil carbon storage. In essence, understanding the complex interplay between lipids in root exudates and soil microbial communities provides valuable insights for sustainable agricultural practices and biofeedstock ecosystem management.

## Methods

### Plant harvest and exudate collection

Five mature tall wheatgrass (*Thinopyrum ponticum*, cultivar: Alkar) plant bunches were harvested in October 2022 from our Tall Wheatgrass Field Trial site at Washington State University (WSU)-Irrigated Agriculture Research and Extension Center (WSU-IAREC) in Prosser, WA, USA. These plants were established in 2018 and were allowed to grow for ∼4 years under normal irrigation conditions. Plants were harvested along with their intact roots and the surrounding soil using a shovel to unearth a root ball that was approximately 30 cm wide x 30 cm deep. After gentle removal of root-associated soil, plant roots were washed in MilliQ water by submerging and shaking gently to remove any remaining soil. After washing, the plants were transferred to 5 L glass beakers where their roots were submerged in 1.5 L of pure milliQ water for 2 hours. The aqueous exudate solutions were vacuum filtered at 0.22 μm and aliquoted into 50 mL tubes for storage at -80 C. Subsequently, the plants roots were transferred to 5 L glass beakers where their roots were submerged in 500 mL chloroform for 5 minutes to collect hydrophobic lipids in exudates. The chloroform exudates were aliquoted into 50 mL tubes for storage at -80 C. 25 mL of aqueous or chloroform exudates were vacuum dried and reconstituted for downstream metabolomics and lipidomics analysis respectively.

Root samples were lyophilized prior to lipid extractions using a modified Folch extraction known as MPLEx (Nakayasu *et al*., 2016, Nicora *et al*., 2018). Lyophilized root samples were ground into a fine powder using a GenoGrinder and 200 mg of powdered sample was used for extraction. Briefly, 5 mL of cold 2:1 (v/v) mixture of chloroform: methanol was added to each sample followed by vortexing. Then 1 mL of cold Milli-Q water was added and samples were vortexed for 1 min. Samples were allowed to cool on ice followed by addition of 1 mL of cold Milli-Q water and vortexing for 1 min. Samples were centrifuged at 4,000 g for 5 min at 4°C resulting in separation into three defined upper layers, the upper-most aqueous (metabolite) layer, the protein interlayer and the lower organic (lipid) layer with the remaining debris at the bottom. The polar metabolite and lipid fractions were collected separately and dried down.

### Mass spectrometry (LC-MS/MS) analysis

Metabolites were analyzed using reverse phase (RP) and hydrophilic interaction chromatography (HILIC) separations on a Thermo Fisher Scientific HF-X mass spectrometer (Thermo Scientific, San Jose, CA) coupled with a Waters Acquity UPLC H class liquid chromatography system (Waters Corp., Milford, MA). Metabolites were brought up in 200 uL of 50% LCMS grade methanol and 50% nanopure water. RP separation was performed by injecting 5 uL of sample onto a Thermo Scientific Hypersil GOLD C18 column (3 µm, 2.1 mm ID X 150 mm L) heated to 40ºC. Metabolites were separated using an 18 minute gradient with data collected on the first 17 minutes. The RP mobile phase A consisted of 0.1% formic acid in nanopure water and a mobile phase B consisting of 0.1% formic acid in LCMS grade acetonitrile. The gradient used was as follows (min, flowrate in mL/min, %B): 0,0.4,10; 2,0.4,10; 11,0.4,90; 12,0.4,90; 12.5,0.5,90; 13.5,0.5,10; 14,0.5,10; 14.5,0.4,10; 15,0.4,10. Positive and negative ion modes were analyzed in separate injections. HILIC separation was performed by injecting 3 uL of sample onto a Waters Acquity BEH Amide column (130 Å, 1.7 µm, 2.1 mm ID X 100 mm L) heated to 55ºC. Metabolites were separated using a 16 minute gradient with data collected for the first 15 minutes. The HILIC mobile phase A consisted of 0.05% ammonium hydroxide, 5% LCMS grade acetonitrile, and 94.95% 10 mM ammonium acetate in nanopure water with a mobile phase B consisting of 0.05% ammonium hydroxide in 99.95% LCMS grade acetonitrile. The gradient used was as follows (min, flowrate in mL/min, %B): 0,0.3,95; 6,0.3,37; 7,0.3,37; 7.1,0.3,95; 7.2,0.5,95; 9.5,0.5,95; 9.7,0.3,95; 16,0.3,95.

Positive and negative ion modes were analyzed in separate injections. For both RP and HILIC separations the Thermo Fisher Scientific HF-X was equipped with a HESI source and high flow needle with the following parameters: spray voltage of 3.6 kV in positive mode and 3kV in negative mode, capillary temperature of 300ºC, probe heater temp of 370ºC, sheath gas at 60 L/min, auxiliary gas at 25 L/min, and spare gas at 2 L/min. Metabolites were analyzed at a resolution of 60 k and a scan range of 50 to 750 m/z for parent ions followed by MS/MS HCD fragmentation which is data dependent on the top 4 ions with a resolution of 15 K and stepped normalized collision energies of 20, 30, and 40. Total organic carbon (TOC) and total nitrogen (TN) in metabolite extracts were measured in acidified extracts on a Vario TOC cube (Elementar, Germany). C/N ratio was calculated by the ratio of TOC and TN.

Lipid extracts were analyzed by RP separation using a Waters Aquity UPLC H class system (Waters Corp., Milford, MA) coupled with a LTQ Velos Elite Orbitrap mass spectrometer (Thermo Scientific, San Jose, CA). Samples were stored in 2 parts chloroform and 1 part methanol, were evaporated, and reconstituted in 100 µL of 90% methanol and 10% chloroform prior to injecting 10 µl onto a Waters column (CSH 3.0 mm x 150 mm x 1.7 µm particle size) heated to 42ºC. Lipids were then separated using a 42 min gradient with data collected on the first 34 minutes. Mobile phase A consisted of 10mM ammonium acetate in 60% nanopure water and 40% LCMS grade acetonitrile with a mobile phase B consisting of 10 mM ammonium acetate in 90% LCMS grade isopropanol and 10% LCMS grade acetonitrile. The gradient used for both positive and negative mode was as follows (min, flowrate, %B): 0,0.25,40; 2,0.25,50; 3,0.25,60; 12,0.25,70; 15,0.25,75; 17,0.25,78, 19,0.25,85; 22,0.25,92; 25,0.25,99; 34,0.25,99; 34.5,0.3,99; 35,0.3,99; 35.5,0.3,99; 36,0.35,40; 37,0.3,40; 38,0.25,40. A Thermo HESI source was coupled to the mass spectrometer inlet with both heated to 350ºC with a spray voltage of 3.5 kV and sheath, auxiliary, and sweep gas flows of 45, 30, and 2, respectively. Positive and negative polarities were analyzed in separate sample runs. Lipids were fragmented by both HCD and CID using a parent ion scan of 200-2000 m/z with a 60 k resolving power. Subsequent MS/MS was performed on the top 4 ions. An isolation width of 2 m/z units and a maximum charge state of 2 were used for both CID and HCD scans. A normalized collision energy of 35 was used for CID while 30 was used for HCD. CID spectra were acquired in the ion trap using an activation Q value of 0.18, while HCD spectra were acquired in the Orbitrap at a mass resolution of 7.5 k and a first fixed mass of m/z 90. For quantification of glycerolipids, a sample was spiked with EquiSPLASH lipid mix (Avanti Polar Lipids) containing deuterated TG, DG and MG internal standards and run in positive and negative mode with each injection containing 0.1 µg of each standard. The area under the peak for the standards was used for quantifying the total amount of lipids of the same lipid class in the samples. Total carbon (TC) and nitrogen (TN) content in lipid extracts was quantified after drying down extracts in tin capsules by using an Elementar Vario Isotope cube elemental analyzer (Elementar, Germany). C/N ratio was calculated by the ratio of TC and TN. Chloroform blanks were analyzed alongside samples to control for residual C contributed by the solvent.

Data processing: Metabolite and lipid identifications were made using MS-DIAL v4.92 for peak detection, identification and alignment. For metabolomics, the experimental data was matched both to in-house libraries (m/z less than 0.003 Da, retention time less than 0.3 min, MS/MS spectral match) and a compilation of publicly available MS/MS databases (m/z less than 0.003 Da, MS/MS spectral match) (VS17) available in the MS-DIAL metabolomics MSP spectral kit (http://prime.psc.riken.jp/compms/msdial/main.html#MSP). The tandem mass spectra and corresponding fragment ions, mass measurement error, and aligned chromatographic features were manually examined to remove false positives. Relative quantification was performed by calculating peak areas on extracted ion chromatograms of precursor masses. Features detected in at least three of the five replicates were retained.

Although confident structural identification of features remains a central goal of metabolomics and lipidomics, it is useful to note that a considerable fraction of features detected in untargeted mass-spectrometry based analyses can arise due to artifacts such as in-source fragments, adducts, isotopes, and contaminants and may not all be biological, unique or informative (Peisl *et al*., 2018). To prepare metabolomics data for CANOPUS analysis, Proteowizard (Chambers *et al*., 2012) and MZmine3 (Schmid *et al*., 2023) were used on each mode separately. Raw data was centroided, peak-picked, and converted to compatible formats with Proteowizard. Resulting mzML files were imported into MZmine3 and underwent noise cancellation and automated data analysis pipeline (ADAP) chromatogram generation. Noise intensity levels were set at 6.0E3–8.0E3 a.u. (arbitrary units) for MS2 spectra and 6.0E4–1.0E5 a.u. for MS1. The local minimum resolver was manually adjusted to reduce peak broadening and splitting; only significant peaks in the chromatograms were recognized as features. Minimum absolute peak height was set to 1.5E5– 2.5E5 a.u. across the modes. 13C related peaks within the same feature list were removed such that only the most significant isotope remained. The isotope pattern finder was also used to detect stable isotopes of hydrogen, carbon, nitrogen, oxygen, sulfur, and phosphorus. Deisotoped feature lists were aligned to produce one feature list with data from the entire mode. Further analysis filtered out rows with less than two features present and remaining ^13^C isotopes. Finally, as feature detection steps (e.g. peak height below minimum intensity, alignment errors, etc.) may result in lost peaks, gap filling was performed. The resulting aligned, filtered, and gap-filled feature list of each mode was exported as an MGF file. Molecular networking files were saved for downstream analysis. A total of 1559 high quality features with MS2 scans were obtained across the 4 modes.

MGF files were transferred to SIRIUS 5.7.2. (Duhrkop *et al*., 2019) for CANOPUS analysis. CANOPUS utilized the fragmentation trees and molecular fingerprints to predict ClassyFire classifications of features (Djoumbou Feunang *et al*., 2016, Duhrkop *et al*., 2021). Classifications with certainties/confidence below 75% were discarded. Summaries were exported with CANOPUS ClassyFire predictions.

## Funding and Acknowledgements

PNNL is a multi-program national laboratory operated by Battelle for the DOE under Contract DE-AC05-76RLO 1830. This program is supported by the U. S. Department of Energy, Office of Science, through the Genomic Science Program, Office of Biological and Environmental Research, under FWP 70880. A portion of this work was performed in the William R. Wiley Environmental Molecular Sciences Laboratory (EMSL), a national scientific user facility sponsored by the Office of Biological and Environmental Research and located at PNNL. IY and DH were supported by the DOE Office of Science’s Science Undergraduate Laboratory Internship (SULI) program. The authors thank Jayde Aufrecht, Daisy Herrera, Sharon Zhao, Eric Garza, Meagan Burnet and Jesse Trejo for assisting with root exudate sample collection following the plant harvest, Sophia McKever and Kaizad Patel for assistance with TC/TN measurements and Shannon Sheridan for helping curate the data package on DataHub.

## Author Contributions

SPC conceived the initial idea and SPC and KSH designed the study. SPC and SLB carried out field collection of plants and exudates. SLB measured TC and TN content in samples. JE collected the LC-MS/MS data on the instrument and helped with glycerolipid quantification. SPC and IY analyzed the metabolomics and lipidomics data and wrote the initial draft of the manuscript. DH analyzed the root tissue lipidomics data. All authors contributed to produce the final draft. KSH obtained the funding.

## Data Availability

Primary raw mass spectrometry lipidomics and metabolomics data have been deposited at MassIVE under the accession: MSV000093600 (will be made public after peer-review when manuscript is accepted for publication).

